# Antimicrobial resistance, pathogenicity and virulence patterns of *Escherichia coli* isolated from cockroaches (*Blattella germanica*) across diverse environments: a public health concern

**DOI:** 10.1101/2024.12.29.630637

**Authors:** Sanzila Hossain Sigma, Md. Nahid Ashraf, Susmita Karmakar, Sabrina Sultana Rimi, Sourav Chakraborty, Mahbubul Pratik Siddique, Jayedul Hassan, Md. Alimul Islam, Md. Tanvir Rahman, Md. Safiqul Islam, Muhammad Tofazzal Hossain

## Abstract

This investigation explores the prevalence and antimicrobial resistance (AMR) of *Escherichia coli* isolated from *Blattella germanica* (cockroaches) from diverse locations, including homes, kitchens, and laboratories, over the course of six months, from August 2022 to January 2023. A total of 125 cockroaches were analyzed, yielding 67 (53%) of positive *E. coli* isolates, with kitchen environments having the highest incidence (57.77%). The existence of virulence genes (*stx1*, *stx2*, and *rfbO157*) was confirmed by pathogenicity assessments carried out on mouse model, which led to a considerable increase in morbidity and mortality. 82.08% of isolates showed evidence of resistance to at least one antibiotic, according to the antimicrobial susceptibility test with β-lactams displaying the highest rates of resistance. Remarkably, complex resistance patterns were observed in 77.61% of the isolates, which were categorized as multidrug-resistant. Multiple antibiotic resistance genes (ARGs) were found by molecular analysis, particularly *bla_TEM_* and *tetA* as well as virulence-associated genes (VAGs) linked to extraintestinal pathogenic *E. coli*. Phylogenetic grouping indicated that 90.38% of the MDR isolates belonged to virulent groups B2 and D. These findings highlight the role of cockroaches as potential reservoirs for pathogenic *E. coli*, raising significant public health concerns regarding AMR. The study underscores the imperative need for thorough investigation and feasible control strategies to ease the dissemination of AMR in diverse contexts.

## Introduction

Cockroaches are abounded globally; over 4,000 species of cockroaches have been documented yet. Due to their strong synanthropic association with human environments, particularly food preparation areas, they pose a severe risk to public health [1–3]. These pests are most at home in filthy areas with easy access to waste and food, like sewage systems, latrines, rubbish dumps, and poultry buildings [4]. Cockroaches have been found to harbor and transfer pathogenic microorganisms through their cuticles or gut, resulting in the contamination of food and surfaces during their foraging activities [5–8]. Their proclivity towards dark, warm, and humid settings as well as their ability to endure in unhygienic conditions make them perfect transferors of pathogenic microbes [4,9,10].

Research has indicated that cockroaches can carry up to 32 different pathogenic bacteria on their surfaces and in their feces, when they travel through diverse settings [11]. According to estimations, they may have up to 14 million organisms on their body parts and 7 million in each feces drop, which is a worrisome level of bacterial load [12]. The predominant bacterial species recovered from cockroaches are Gram-negative, particularly those belonging to the *Enterobacteriaceae* family [13,14]. Cockroaches have been implicated in outbreaks of dysentery, with multiple enteropathogenic bacteria as well as toxigenic strains of *Escherichia coli* remaining viable in their gut for several days [15–17]. Crucially, cockroaches exacerbate public health attempts to control disease transmission by aiding in the expand of antibiotic-resistant bacteria [18–20].

AMR or resistance to antimicrobial substances is a key comprehensive health matter, since it reduces the potency of treatments for infections, resulting in increased mortality, longer hospitalizations, and higher healthcare expenses. MDR organisms are especially worrisome because they restrict treatment options and raise the likelihood of illness that are challenging or impossible to medicate [21,22].

*E. coli* is a commensal microbiota found in the GI tract of all warm-blooded mammals in general [23,24]. Some strains of the bacteria have attained pathogenicity by acquiring virulence genes or resistance through horizontal gene transfer, which can result in a variety of intestinal and extra-intestinal diseases. A cause for serious foodborne diseases, hemorrhagic colitis, and hemolytic uremic syndrome, *Escherichia coli O157:H7* and STEC isolated from cockroaches are of special concern [14,25–27]. With as little as 10–100 cells, this strain can infect people, and its potent Shiga toxins have been connected to epidemics across the globe [28]. In Bangladesh, *E. coli*, especially enterotoxigenic strains (ETEC) is a prominent source of childhood diarrhea, accounting for up to 20% of total cases [29]. Recent study have provided evidence of cockroaches harboring pathogenic bacteria and associated resistance patterns in Bangladesh with highest prevalence of *E .coli* [13]. Hence, the role of cockroaches in spreading pathogenic and multidrug-resistant (MDR) *E. coli* in Bangladesh’s tropical climate is an area of growing concern.

Despite the known risks, comprehensive studies on cockroaches as carriers of pathogenic and MDR *E. coli*, in Bangladesh remain limited. This study aims to bridge that gap by isolating *E. coli* from cockroaches and conducting a detailed analysis of their phylogenetic grouping, virulence genes, pathotypes, and antibiotic resistance patterns. By doing so, it will shed light on the critical public health implications of these urban pests and their potential to disseminate antibiotic-resistant infections.

## Methodology

### Area of study and data collection

125 cockroaches in total were collected from August 2022 to January 2023 from kitchen, washroom, premises of household and BAU residential halls and Department of Microbiology and Hygiene. Collected cockroaches were transported to microbiology laboratory in sterile bags. Details on samples are stated in **S1 Table in S1 File.**

### Isolation of bacteria

Chilling cockroaches at 0°C for five to ten minutes rendered them immobile. Using sterile dissecting equipment, aseptic dissection of the mouth, legs, and gut (internal component) was carried out. The samples were then put in different test tubes that contained nutrient broth. A single loop of the broth was collected and streaked onto an EMB agar plate (HiMedia, India) following an overnight enrichment period. After that, the streaked Petri dishes were incubated at 37°C overnight. Different morphological properties, including as colony characteristics on EMB agar, Gram staining features, and biochemical assays such sugar fermentation, the indole test, and the methyl red test, were used to identify between *Escherichia coli* isolates [30].

### Molecular identification of *E. coli*

Polymerase chain reaction was applied to confirm *E. coli* by using primers designed against the *malB* gene (**Table 1**). Following the preceding formulation of Islam et al., 2021 [31], the PCR procedure started with the extraction of chromosomal DNA from isolated *E. coli* with the boiling principles.

**Table1.**
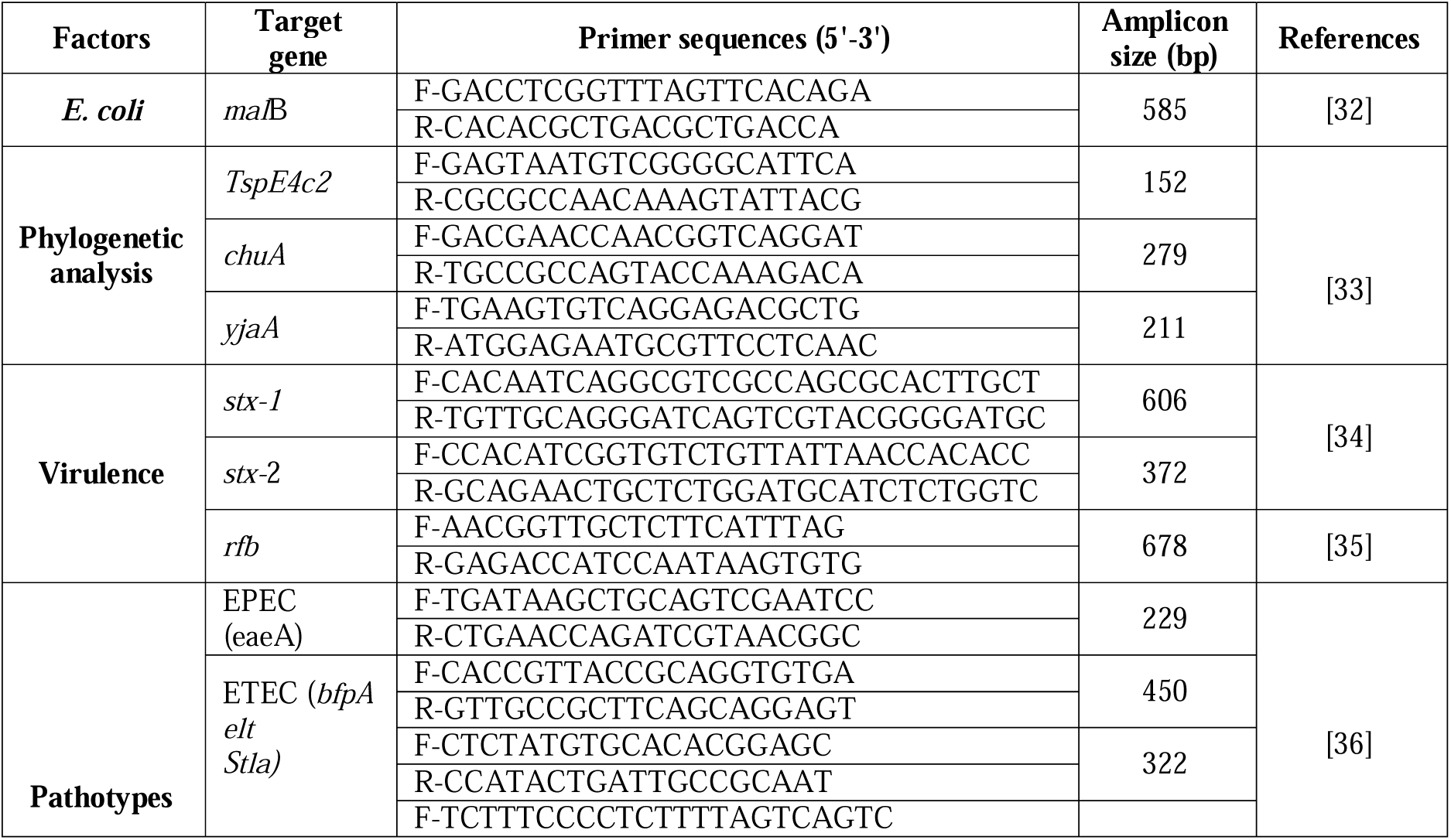
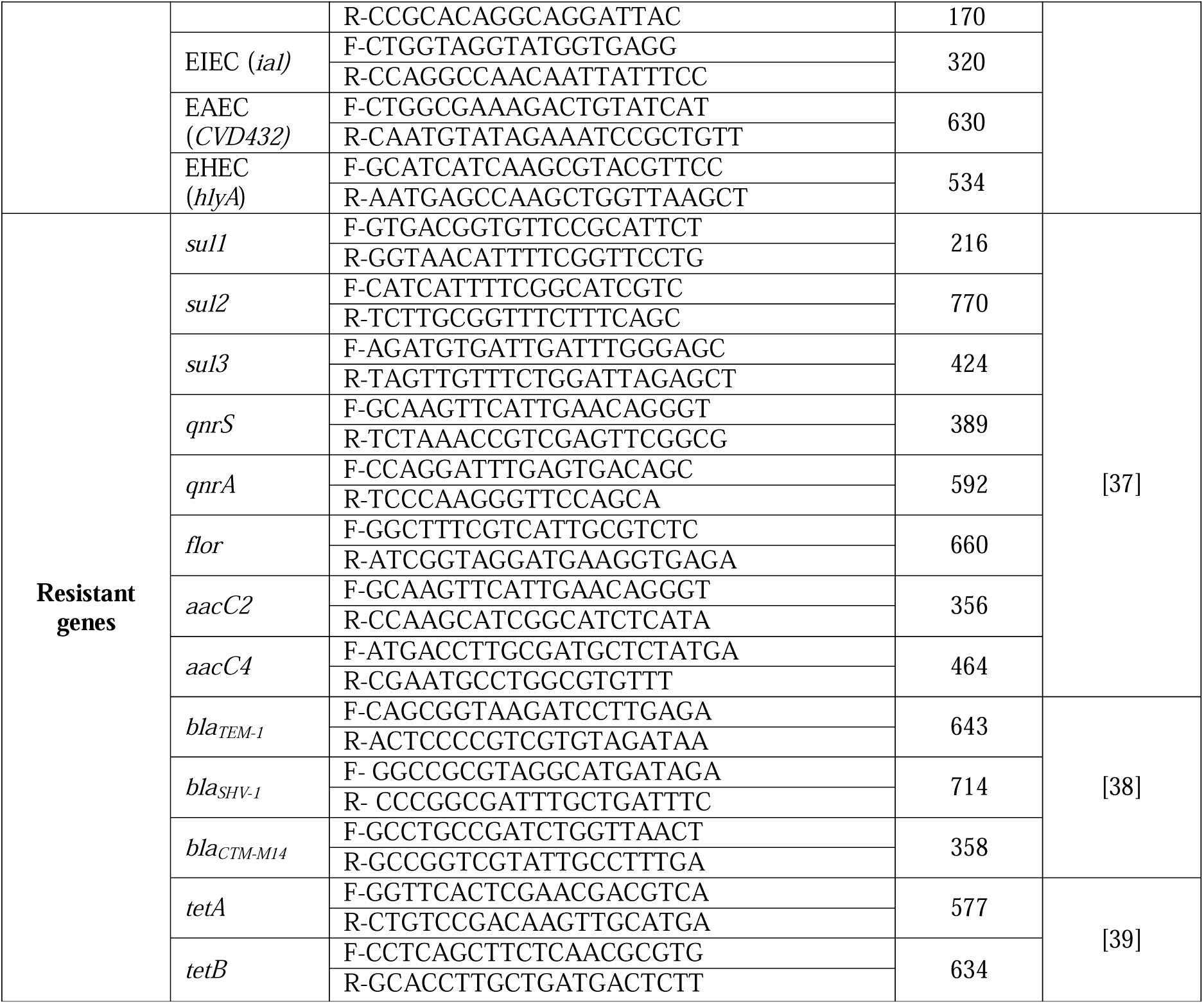
List of primers used in the study.

### Phylogenetic classification of *E. coli*

*E. coli* isolates were analyzed for phylogenetic grouping (A/B1/B2/D) using PCR pointing *chuA*, *yjaA*, and *TSPE4.C2* in accordance with previous procedure [33].

### Virulence associated genes (VAGs) identification in *E. coli*

By using of three pairs of primers designed from the VAGs often present in *E. coli*, such as *stx1*, *stx2*, and *rfbO157* the genes were differentiated using duplex PCR. **Table 1** provides an overview of the primer sequences as well as relevant references.

### Pathotypes identification in *E. coli*

The Hegde et al. (2012) approach was followed in order to identify the five types of diarrheagenic *E. coli*: enteropathogenic *E. coli* (EPEC), enterohemorrhagic *E. coli* (EHEC), enterotoxigenic *E. coli* (ETEC), enteroaggregative *E. coli* (EAEC), and enteroinvasive *E. coli* (EIEC) [36]. Specific primers indicated in **Table 1** were used for multiplex PCR.

### Pathogenicity test

The pathogenicity test was also performed in mice model [AWEEC/BAU/2018(28)] via oral administration of *E. coli* strains of *stx1, stx2* and *rfbO157* following Wadolkowski [40]. Pathogenic *E. coli* bacteria isolation and identification from intestine of mice were also studied by culture and PCR method.

### Antibiotic susceptibility test (AST)

AST of *E. coli* isolates was executed as stated by the Kirby-Bauer disc diffusion method. Freshly cultured bacteria, adjusted to 0.5 McFarland standards, were spread on Muller-Hinton agar (HiMedia, India) [41]. Sixteen commonly used antibiotics from seven categories were selected for the study: aminoglycosides (amikacin-30µg, gentamicin-10μg, streptomycin-10μg), tetracyclines (tetracycline-30μg, doxycycline-30μg), β-lactams [cephalosporins (cefixime-30μg, cefalexin), carbapenems (imipenem-10μg), penicillins (ampicillin-10μg, amoxicillin-clavulanic acid-30µg)], macrolides (azithromycin, erythromycin-15µg), quinolones (ciprofloxacin-5μg, levofloxacin-5µg), amphenicols (chloramphenicol-30µg), and sulfonamides (trimethoprim-sulfamethoxazole, SXT, 10μg). To assure accuracy, the experiment was conducted at least three times. The *E. coli* isolates’ antibiotic susceptibility profiles (sensitive, intermediate, and resistant) were evaluated according to the Clinical and Laboratory Standards Institute’s 2022 criteria [42]. Isolates exhibiting resistance to three or more antibiotic classes were classified as multidrug-resistant (MDR) according to Magiorakos [43]. MAR = u/v is the formula that was used to assess the multiple antibiotic resistance (MAR) index, where “u” symbolizes the number of antibiotics to which an isolate was resistant and “v” symbolizes the total number of antibiotics examined in this investigation [44].

### Detection of Antibiotic-resistant genes (ARGs)

Primers of 13 AGRs in 6 categories were used to detect the AGRs from isolates. The PCR primer sequences, amplicon size with references were depicted in **Table 1**.

## Results

### Prevalence and population density of cockroaches

A total of 125 cockroaches were captured and trapped and *Blatella germanica* were detected in surveyed locations. Among the captured places, cockroaches had the highest dominant in residential halls 71 (56.8%) and kitchen areas 45 (36.0%) compared with other captured places. Moreover, regarding distinct phases of life, 37, 88 nymph and adults were detected respectively. Both phases were more ubiquitous in kitchen settings (**S1 Table in S1 File**).

### Occurrence of *E. coli*

Out of 125 cockroaches, 67 (53%) were positive for *E. coli* and detection rate was higher in kitchen areas 26 (57.77%). In distinct life phases, the detection rate of *E. coli* in cockroach, adults were higher 52 (59.09%) than that of nymphs. Additionally, isolation rate was higher from the gut (32.80%) compared to external surface of cockroach (**S2 Table in S1 File, Fig 1**)

**Fig 1.** Occurrence of *E. coli* in relation to (a) Life stage and body parts (b) Different places and areas.

### Antimicrobial resistance (AMR) evaluation of 67 isolates and profiling of MDR strains

Out of 67 *E. coli* isolates, 55 strains (82.08%, 55/67) showed resistance to at least one antimicrobial agent, while only 12 strains (17.91%, 12/67) were sensitive to all agents (**Table 2**, **Fig. 2**). Resistance to β-lactam antibiotics was the highest, with 82.08% (55/67) of strains affected, and followed by tetracycline at 71.64% (48/67) and sulfonamides at 67.16% (45/67). The lowest resistance rate was observed for macrolides, at 34.33% (23/67). Specifically, resistance to AMP (82.08%, 55/67), TET (71.64%, 48/67), and SXT (67.16%, 45/67) ranked highest among the 16 tested antimicrobial agents, while AK had the lowest resistance rate at 14.93% (10/67). The resistance rates for the remaining 12 antibiotics ranged from 17.91% (IPM) to 52.24% (C). Further analysis of resistance phenotypes revealed that 77.61% (52/67) of the isolates were classified as MDR *E. coli* (**Fig. 2**). Among these 52 MDR strains, 24 distinct resistance patterns were witnessed. The most common patterns included AMC/AMP/AZM/SXT/C/CFM/CN/DO/E/IMP/TE/S/GEN,AMC/AMP/AZM/CFM/C/SXT/DO/S/T E/LEV/GEN,AMP/AMC/AZM/CIP/SXT/DO/E/CN/LEV/TE/AK/S,AMP/C/SXT/AMC/CFM/E/IPM/TE/GEN/C and AMP/AMC/C/CFM/TE/S/LEV/GEN. Remarkably, the MDR isolates showed resistance to 3 to 7 different antibiotic categories, with MAR indices ranging from 0.2 to 0.8 (**Table 3**).

**Table 2.**
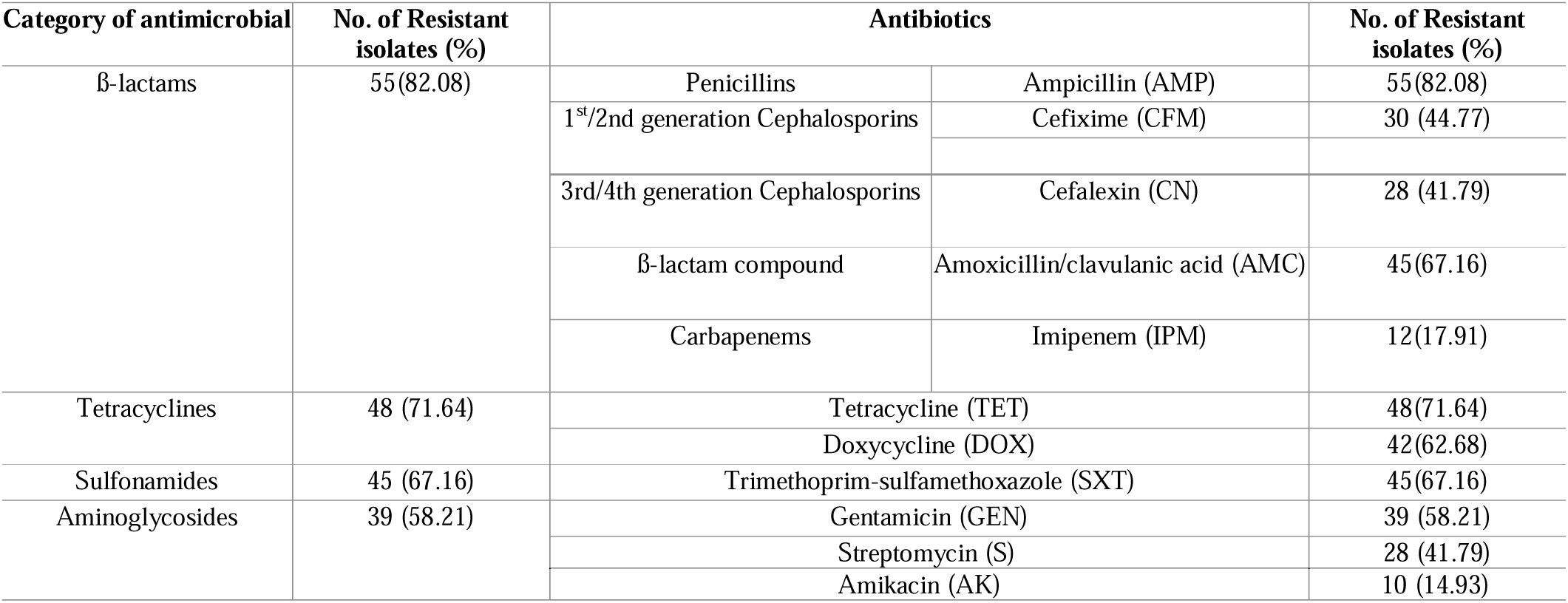
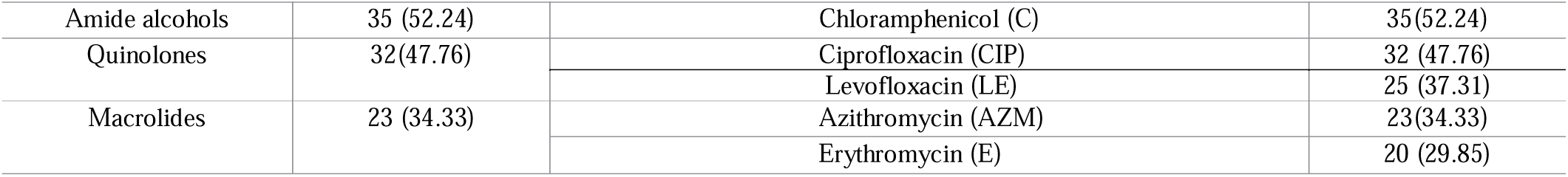
Antimicrobial resistance (AMR) detected in *E. coli* strains isolated from cockroaches (n = 55).

| Category of antimicrobial | No. of Resistant isolates (%) | Antibiotics |  | No. of Resistant isolates (%) |
| --- | --- | --- | --- | --- |
| β-lactams | 55(82.08) | Penicillins | Ampicillin (AMP) | 55(82.08) |
|  |  | 1<sup>st</sup>/2nd generation Cephalosporins | Cefixime (CFM) | 30 (44.77) |
|  |  | 3rd/4th generation Cephalosporins | Cefalexin (CN) | 28 (41.79) |
| | | β-lactam compound | Amoxicillin/clavulanic acid (AMC) | 45(67.16) |
|  |  | Carbapenems | Imipenem (IPM) | 12(17.91) |
| Tetracyclines | 48 (71.64) | Tetracycline (TET) |  | 48(71.64) |
|  |  | Doxycycline (DOX) |  | 42(62.68) |
| Sulfonamides | 45 (67.16) | Trimethoprim-sulfamethoxazole (SXT) |  | 45(67.16) |
| Aminoglycosides | 39 (58.21) | Gentamicin (GEN) |  | 39 (58.21) |
|  |  | Streptomycin (S) |  | 28 (41.79) |
|  |  | Amikacin (AK) |  | 10 (14.93) |

|  |  |  |  |
| --- | --- | --- | --- |
| Amide alcohols | 35 (52.24) | Chloramphenicol (C) | 35(52.24) |
| Quinolones | 32(47.76) | Ciprofloxacin (CIP) | 32 (47.76) |
|  |  | Levofloxacin (LE) | 25 (37.31) |
| Macrolides | 23 (34.33) | Azithromycin (AZM) | 23(34.33) |
|  |  | Erythromycin (E) | 20 (29.85) |

**Fig 2.** (a) 0R symbolizes the isolates which were sensitive to used antibiotics and 1-7R symbolizes isolates which exhibited resistant to 1 to 7 categories of antibiotics. (b) Highlighted bars depict the phenotypic resistance patterns (occurring 3 times) in 52 MDR isolates with number MDR strains. Dominate resistance patterns are highlighted by the red boxes.

**Table 3.** Antibiotic Resistant pattern and MAR index.

| No. of patterns | Antibiotic Resistant pattern | No. of antibiotic (Classes) | No. of isolates | Overall MDR isolates | MAR index |
| --- | --- | --- | --- | --- | --- |
| 1 | AMP, AMC, C, CFM, GEN, CIP, E, CN, IMP, TE, DO, S, SXT, LEV | 13(7) | 3 | 52/67<br>(77.61) | 0.8 |
| 2 | AMP, AMC, AZM, CIP, SXT, DO, E, CN, LEV, TE, AK, S, C | 13(7) | 5 |  | 0.8 |
| 3 | AMC, AMP, AZM, SXT, C, CFM, CN, DO, E, IMP, TE, S, GEN | 13 (5) | 7 |  | 0.8 |
| 4 | AMC, AMP, AZM, CFM, C, SXT, DO, S, TE, LEV, GEN | 11(7) | 6 |  | 0.7 |
| 5 | AMP, C, SXT, AMC, CFM, E, IPM, TE, GEN | 9(6) | 3 |  | 0.6 |
| 6 | AMC, AMP, AZM, DO, E, AK, S, TE, GEN | 9(5) | 2 |  | 0.6 |
| 7 | AMP, AMC, C, CFM, TE, S, LEV, GEN | 8(5) | 5 |  | 0.5 |
| 8 | AMP, AMC, AZM, IPM, SXT, C, GEN | 7(5) | 2 |  | 0.4 |
| 9 | AMP, CFM, DO, TE, SXT, LEV, GEN | 7(5) | 1 |  | 0.4 |
| 10 | AMP, AMC, CN, TE, AK, SXT, C | 7(5) | 2 |  | 0.4 |
| 11 | AMC, AMP, E, SXT, DO, TE, C | 7(5) | 1 |  | 0.4 |
| 12 | AMP, C, DO, TE, SXT, GEN | 6(5) | 2 |  | 0.4 |
| 13 | AMC, AMP, CFM, AZM, TE, SXT | 6(4) | 1 |  | 0.4 |
| 14 | AMP, DO, LEV, AK, TE, SXT | 6(5) | 1 |  | 0.4 |
| 15 | AMC, AMP, C, TE, SXT, LEV | 6(5) | 1 |  | 0.4 |
| 16 | AMC, AMP, C, CFM, TE, GEN | 6(4) | 1 |  | 0.4 |
| 17 | AMC, AMP, CN, LEV, TE, SXT | 6(4) | 1 |  | 0.4 |
| 18 | AMC, AMP, C, DO, TE, SXT | 6(4) | 1 |  | 0.4 |
| 19 | AMP, SXT, C, DO, TE, GEN | 6(5) | 1 |  | 0.3 |
| 20 | AMC, AMP, DO, IPM, TE, C | 6(3) | 1 |  | 0.4 |
| 21 | AMP, CFM, SXT, TE, LEV | 5(4) | 1 |  | 0.3 |
| 22 | AMP, DO, TE, SXT, GEN | 5(4) | 2 |  | 0.3 |
| 23 | AMC, AMP, SXT, DO, TE | 5(3) | 1 |  | 0.3 |
| 24 | AMP, SXT, TE | 3(3) | 1 |  | 0.2 |

AMC = Amoxyclave, AMP = Ampicillin, AZM = Azithromycin, AK = Amikacin, SXT = Trimethoprim-sulfamethoxazole, CN = Cefalexin, CFM = Cefixime, CIP = Ciprofloxacin, DOX = Doxycycline, E = Erythromycin, IPM = Imipenem, LEV = Levofloxacin, C = Chloramphenicol, GEN = Gentamycin, S = Streptomycin, TET = Tetracycline.

### Phylogenetic grouping of MDR *E. coli* isolates

Based on phylogenetic study investigations, group B2 was found to be the one with the highest prevalence, making up 67.30% (35/52) of the 52 MDR *E. coli* strains. Group D came in next with 23.07% (12/52) and group B1 with 9.61% (5/52). Remarkably, only 9.46% of the MDR *E. coli* strains belonged to the commensal group B1, whereas 90.38% (47/52) were linked to the virulent extraintestinal-related groups B2 and D.

### Occurrence and dissemination of antibiotic resistance genes (ARGs) and virulence-associated genes (VAGs) in multidrug-resistant (MDR)

#### E. coli strains

Eleven of the eighteen ARGs across six categories were detected (**Table 4**). ARGs *tetA* and *bla_TEM_*exhibited the highest detection rates, both at 86.54% (45/52) in the study. Additionally, *sul2*, *qnrS*, and *flor* had detection rates exceeding 65.38%, 59.61%, and 51.92%, respectively. The remaining ARGs had detection rates ranging from 48.07% (*bla_SHV_*) to 23.07% (*tetB*). Subsequent investigation revealed one variant for the *bla_TEM_* and *bla_SHV_* genes, with detection rates of 86.54% (45/52) for *bla_TEM-1_* and 48.07% (25/52) for *bla*_SHV-1_. Among virulence-associated genes (VAGs), *rfb0157* had the highest detection rate at 61.54% (32/52), followed by *stx1* at 48.07% (27/52) and *stx2* at 23.07% (12/52). Analysis of five diarrheagenic *E. coli* (DEC) pathotypes revealed that ETEC, EAEC, and EHEC, which are defined by the presence of virulence genes like *elt*, *stlA*, *hlyA*, and *eaeA* were detected. Among these 3 pathotypes, ETEC was the most ubiquitous (57.69%) (**S3 Table in S1 File**, **Fig. 3**).

**Table 4.** Antimicrobial resistance (AMR) and Antibiotic-resistant genes (ARGs) detected in MDR *E. coli* (n=52)

| Category of antimicrobial | No. of Resistant Isolates (%) | Antibiotics | No. of Resistant Isolates (%) | ARGs | No. of Positive Isolates (%) |
| --- | --- | --- | --- | --- | --- |
| β-lactams | 52 (100) | AMP | 52 (100) | *bla<sub>TEM</sub>* | 45(86.54) |
|  |  | CFM | 25 (48.08) |  |  |
|  |  | CN | 23 (44.23) |  |  |
|  |  | AMC | 43 (82.69) | *bla<sub>SHV</sub>* | 25(48.07) |
|  |  | IPM | 8 (15.38) |  |  |
| Tetracyclines | 45 (86.54) | TET | 45 (86.54) | *tetA* | 45(86.54) |
|  |  | DOX | 38 (73.07) | *tetB* | 12(23.07) |
| Sulfonamides | 40 (76.92) | SXT | 40 (76.92) | *sul2* | 34(65.38) |
|  |  |  |  | *sul1* | 20(38.46) |
|  |  |  |  | *sul3* | 13(25.00) |
| Aminoglycosides | 37(71.15) | GEN | 37 (71.15) | *aacC2* | 25(48.07) |
|  |  | S | 22 (42.31) | *aacC4* | 18(34.61) |
|  |  | AK | 5 (9.6) |  |  |
| Amide alcohols | 30(54.05) | C | 30 (54.05) | *flor* | 27 (51.92) |
| Quinolones | 28 (45.95) | CIP | 28(45.95) | *qnrS* | 31(59.61) |
|  |  | LE | 20(38.46) |  |  |

**Fig 3.** Prevalence of VAGs and pathotypes in MDR isolates.

#### Pathogenicity test and re-isolation

Pathogenic property of PCR positive strains of *stx1*, *stx2* and *rfbO157* were confirmed by pathogenicity test through randomly selecting 5 isolates from each group. Per os administration of pathogenic *E. coli* in mice induced illness and death of all mice without control during observation period (**Fig 4**). *E. coli* strains of *stx1* and *stx2* by oral administration caused death of mice in the observation period of 4^th^ and 5^th^ days and *E. coli* strains of O157 caused death of mice in the time period of 5^th^ days.

**Fig 4.** Pathogenicity test in mice.

Dead mice were dissected and intestines were collected and processed for re-isolation of pathogenic strains by primary enrichment in nutrient broth and conventional culture on EMB agar and PCR assay also revealed positive results of *E. coli* strains of *stx*1, *stx*2 and *rfb*O157 which specifies that all the 15 isolates by *stx*1, *stx*2 and *rfb*O157 genes were highly pathogenic.

#### Association between AGRs and AMRs or VAGs in *E. coli* strains

The detection rates of antibiotic resistance genes (ARGs) and antimicrobial resistance (AMR) in 52 MDR *E. coli* strains are detailed in **Table 4**. A high prevalence of the β-lactam resistance gene (*bla_TEM_* 86.54%) was observed, with nearly 100% resistance to β-lactam antibiotics (AMP) (52/52).

For tetracyclines, the *tetA* gene was identified in 45 out of 52 MDR strains (86.54%), corresponding to the same proportion of strains exhibiting tetracycline-resistant phenotypes. Regarding sulfonamides, 76.92% (40/52) of the MDR strains were resistant to SXT, the detection rates for the related resistance genes (*sul1*, *sul2*, and *sul3*) ranged from 25.00% (13 of 52 samples) to 65.38% (34 of 52 samples). For aminoglycosides, only two resistance genes (*aacC2* and *aacC4*) were detected, with rates between 48.07% (25/52) and 34.61% (18/52). Phenotypic resistance to aminoglycosides (GEN, S, and AK) varied from 9.6% (5/52) to 71.15% (37/52). The detection rates of other antibiotic resistance phenotypes and their related ARGs fluctuated. These results depicted that only β-lactam and tetracycline-resistant phenotypes generally aligned with their corresponding ARG detection rates, while others were less consistent. We investigated that virulence genes like *stx1*, *stx2*, and *rfbO157* are associated with broad-spectrum antimicrobial resistance. These genes co-occur with key AMR mechanisms, particularly β-lactamases (e.g., *bla_TEM_*, *bla_SHV_*), tetracycline resistance genes (e.g., *tetA*, *tetB*), and sulfonamide resistance genes (e.g., *sul2*, *sul1*). Pathotypes such as ETEC, EAEC, and EHEC, which are defined by the presence of virulence genes like *elt*, *stlA*, *hlyA*, and *eaeA*, exhibited resistance against different antibiotic categories, suggesting a strong association between virulence and resistance in these MDR isolates. The ETEC isolates had high prevalence of ARGs for example *bla_TEM_* (86.54%), *tetA* (86.54%), *sul2* (65.38%), and *qnrS* (59.61% for quinolone resistance. The EHEC isolates also carried the significant resistance genes (*bla_TEM_*, *sul2*, *tetA*, *qnrS*), though the prevalence of *bla_SHV_* was less frequent in this group. ARGs such as *bla_TEM_*, *tetA*, *sul2*, and *qnrS* were also found to be dominant among EAEC isolates.

## Discussion

This study offers critical insights into the distribution, abundanceNo data on counts TCC, and microbial burden of *B. germanica* (German cockroach), with a specific focus on their role as vectors for *Escherichia coli* (*E. coli*) and their resistance to antimicrobial agents. The findings reveal an alarming prevalence of cockroaches in residential and kitchen areas, and the study highlights their potential role in transmitting pathogenic and multidrug-resistant (MDR) bacterial strains. In our study, we analyzed these findings in the context of public health and antimicrobial resistance.

The study demonstrated that *B. germanica* was found in all surveyed locations, with the highest prevalence in residential halls (56.8%) and kitchens (36.0%). Among the most common urban pests is the German cockroach, *Blattella germanica*, which is frequently found in household, commercial and laboratory settings [27,45]. Its widespread presence is attributed to its adaptability to diverse environments, particularly those associated with human habitation, such as kitchens and residential halls [45,46]. This pattern is consistent with other studies that suggest cockroaches thrive in human-inhabited areas with easy access to food and moisture [47]. The higher prevalence of adults (77.6%) compared to nymphs (22.4%) could be attributed to favorable conditions in these environments, allowing for complete development of the insect life cycle.

The detection of *E. coli* in 53.6% of cockroaches highlights their potential role as mechanical vectors for pathogenic bacteria. *E. coli* presence was highest in kitchen areas (57.77%) and washrooms (60.60%), which align with previous research linking cockroaches to contamination in food handling areas [18,48]. Notably, adults had a significantly higher detection rate of *E. coli* (59.09%) compared to nymphs (40.54%), which might be due to their greater mobility and ability to come into contact with contaminated surfaces. This observation aligns with the idea that adult cockroaches are more likely to spread pathogens in their environment [49]. The elevated *E. coli* presence in the gut (32.80%) compared to external body parts (20.80%) indicates that cockroaches can harbor these pathogens internally, possibly through ingestion of contaminated food or waste. The isolation of bacterial isolates from the exterior surface (36.7%) compared to the gut (63.3%) of the cockroach is consistent with earlier investigations [50–52].

The phylogenetic analysis revealed that most MDR *E. coli* strains belonged to group B2 (67.3%) and group D (23.07%), which are associated with extraintestinal pathogenic *E. coli* (ExPEC) [33]. These findings are alarming as ExPEC strains are commonly linked to serious infections such as urinary tract infections and sepsis [53]. The presence of virulent phylogroups in cockroaches raises concerns about their potential to transmit dangerous pathogens in environments shared with humans.

The study’s pathogenicity test using strains positive for *stx1*, *stx2*, and *rfbO157* confirmed the high pathogenicity of *E. coli* strains isolated from cockroaches. The death of all mice within five days following oral administration of these strains highlights the potential public health risk posed by cockroaches as carriers of highly pathogenic *E. coli*. The re-isolation of the strains from the intestines of dead mice further validated the pathogenicity of these isolates, demonstrating their capacity to cause severe illness. The identification of specific (VAGs) in the MDR strains enhances our understanding of the potential health risks linked to cockroach infestations [37].

AMR was prevalent among isolates, with β-lactam resistance being the most common (82.08%). The high resistance to ampicillin (82.08%) and other antibiotics like tetracycline (71.64%) and sulfonamides (67.16%) echoes previous studies documenting the extensive use of these antibiotics in healthcare, agricultural and municipal settings [14,54]. Notably, 77.61% isolates were identified as MDR, and the observed resistance patterns reflected exposure to an extensive range of antibiotics, consistent with global trends in MDR bacterial pathogens [43].

The high prevalence of ARGs, particularly *bla_TEM_* and *tetA*, in MDR strains reflects the strong selective pressure exerted by β-lactam and tetracycline antibiotics. The detection of ExPEC-related virulence-associated genes (VAGs), such as *stx1* (48.07%) and *rfbO157* (61.54%) as well as diarrheagenic pathotypes 57.69% ETEC (*elt,stlA*), 48.07% EAEC (*CVD432*) and 26.92% EHEC (*hlyA,eaeA*) indicates that these strains are not only resistant but also potentially pathogenic to produce serious diseases [36]. This combination of resistance and virulence represents a serious public health concern, as these cockroach-associated *E. coli* strains may be capable of causing severe human diseases [55].

The findings of this investigation reinforce the ability of cockroaches as reservoirs for MDR and potentially pathogenic *E. coli*. Their presence in residential and food preparation areas poses significant health risks, emphasizing the need for comprehensive sanitation and pest control measures combined with robust hygiene practices that cleanliness could help reduce cockroach populations and limit the dissemination of bacteria. Future investigation is requisite to scrutinize the potential for these cockroach-associated *E. coli* strains to spread and cause human infections.

## Conclusions and limitations

This study provides important evidence linking cockroaches to the dissemination of pathogenic and MDR *E. coli* strains, with significant implications for food safety and public health. These findings call for increased attention to pest control, improved sanitation in high-risk areas like kitchens, and continued monitoring of AMR in urban pests. The potential for cockroaches to act as reservoirs of both bacteria and resistance genes highlights the need for coordinated public health strategies to mitigate these risks.

The current study was limited by its sample size and the fact that samples were only collected from the Bangladesh Agricultural University campus. Future research should aim to include a larger sample size and broaden the sampling locations. Additionally, Whole Genome Sequencing could yield more detailed insights into the MDR *E. coli* strains found in cockroaches.

## Funding

Sanzila Hossain Sigma received a National Science and Technology (NST) Fellowship from Ministry of Science and Technology, Government of the People’s Republic of Bangladesh.

## Acknowledgments

The authors would like to thank the authorities for their assistance with this study.

## Competing interests

The authors state no competing interests.

## Author contributions

**Conceptualization:** Muhammad Tofazzal Hossain.

**Data organization:** Sanzila Hossain Sigma.

**Formal analysis:** Sanzila Hossain Sigma, Md. Nahid Ashraf

**Investigation:** Sanzila Hossain Sigma, Susmita Karmakar, Md. Shafiqul Islam, Muhammad Tofazzal Hossain

**Methodology:** Sanzila Hossain Sigma, Md. Nahid Ashraf, Susmita Karmakar, Sabrina Sultana Rimi, Sourav Chakraborty

**Resources:** Md. Shafiqul Islam, Muhammad Tofazzal Hossain, Jayedul Hassan.

**Supervision:** Muhammad Tofazzal Hossain, Md. Shafiqul Islam.

**Validation:** Mahbubul Pratik Siddique, Muhammad Tofazzal Hossain, Md. Shafiqul Islam, Jayedul Hassan.

**Visualization:** Sanzila Hossain Sigma, Muhammad Tofazzal Hossain, Md. Shafiqul Islam, Md. Tanvir Rahman.

**Writing – original draft**: Sanzila Hossain Sigma, Md. Nahid Ashraf

**Writing – review & editing**: Md. Shafiqul Islam, Muhammad Tofazzal Hossain, Md. Tanvir Rahman.

## Notes

### Competing Interest Statement

The authors have declared no competing interest.

